# Efficiency of genome-wide association study in random cross populations

**DOI:** 10.1101/105833

**Authors:** José Marcelo Soriano Viana, Gabriel Borges Mundim, Hélcio Duarte Pereira, Andréa Carla Bastos Andrade, Fabyano Fonseca e Silva

**Author notes:** Corresponding author: José Marcelo Soriano Viana. Federal University of Viçosa, Department of GeneralBiology,36570-900,Viçosa,MG,Brazil. Reference number for data available in public repository:https://dx.doi.org/10.6084/m9.figshare.3201838.version3. REALbreeding* private link: https://figshare.com/s/618bee7accd410464232.

## Abstract

Genome-wide association studies (GWAS) with plant species have employed inbred lines panels. Our objectives were to present additional quantitative genetics theory for GWAS, evaluate the relative efficiency of GWAS in non-inbred and inbred populations and in an inbred lines panel, and assess factors affecting GWAS. Fifty samples of 400 individuals from populations with linkage disequilibrium were simulated. Individuals were genotyped for 10,000 single nucleotide polymorphisms (SNPs) and phenotyped for traits controlled by 10 quantitative trait loci (QTLs) and 90 minor genes, assuming different degrees of dominance and heritabilities of 40 and 80%. The average SNP density was 0.1 centiMorgan and the QTL heritabilities ranged from 3.2 to 11.8%. To increase the QTL detection power, the additive-dominance model must be fitted for traits controlled by dominance effects but must not be fitted for traits showing no dominance. The power of detection was maximized increasing the sample size to 400 and the false discovery rate (FDR) to 5%. The average power of detection for the low, intermediate, and high heritability QTLs were 9.7, 32.7, and 87.7%, respectively. Under sample size of 400 the observed FDR was equal to or lower than the specified level of significance. The association mapping was highly precise. The analysis of the inbred random cross population provided essentially the same results from the non-inbred population. The inbred lines panel provided the best results concerning the low and intermediate heritability QTL detection power, FDR, and mapping precision. The FDR is mainly affected by population structure, compared to relationship information.

## INTRODUCTION

Association mapping is a high-resolution method for mapping quantitative trait locus (QTL) based on linkage disequilibrium (LD) (Yu and Buckler 2006). Linkage disequilibrium is commonly defined as the non-random association of alleles at two loci carried on the same gamete, caused by their shared history of mutation and recombination (Weir 2008). Association mapping has been successful in detecting genes controlling human diseases and quantitative traits in humans and plant and animal species (Pearson and Manolio 2008; Zhu *et al.* 2008; Barendse *et al.* 2007). There are two main association mapping strategies: the candidate gene approach, which focuses on polymorphisms in specific genes that control the traits of interest, and the genome-wide association study (GWAS), which surveys the entire genome for polymorphisms associated with complex traits (Rafalski 2010).

With the advent of high-throughput genotyping and sequencing technologies, breeders have used GWAS to identify genes underlying quantitative trait variation. Compared to QTL mapping, which has precision in the range of 10^5^ to 10^7^ base pairs (Yu and Buckley 2006), the main advantage of GWAS is a more precise identification of candidate genes (Zhu *et al.* 2008). Another advantage is the use of a breeding population instead of one derived by crossing two inbred or pure lines (Flint-Garcia *et al.* 2005). However, as highlighted by Weir (2010), the efficiency of GWAS is considerably affected by relatedness and population structure, which can generate spurious association between unlinked marker and QTL. Rafalski (2010) emphasized that the choices of population (due to the degree of LD and genotypic variation), marker density, and sample size are crucial decisions for achieving greater power of QTL detection. Ingvarsson and Street (2011) discussed the influence of population size, extent of LD, trait heritability (precision of phenotyping), and population structure on GWAS efficiency, highlighting that studies with plant species should greatly increase population size to detect QTLs with lower effect (heritability of 1−2%).

Yu *et al.* (2006) proposed a mixed model approach for GWAS analysis called the Q + K (or QK) method, where Q and K are the population structure and kinship matrices, respectively. This method has provided the best results and greatly has improved the control of both type I and type II error rates compared with other methods. Stich and Melchinger (2009) and Yang *et al.* (2010) compared GWAS methods based on simulated and field data. Based on type I error control and power of QTL detection, they concluded that the mixed model approach using only the kinship matrix (K model) to correct for relatedness was more efficient than the approaches controlling only population structure (Q model) and both population structure and relatedness because the spurious associations could not be completely controlled by population structure. Based on simulated inbred lines panel, Bernardo (2013) demonstrated that his models G and QG were superior to the K and QK, respectively, where G indicates a model that uses genome-wide markers to account for QTLs on background chromosomes. The new approach showed a better balance between power of QTL detection and false discovery rate (FDR). Frąszczak and Szyda (2016) also compared GWAS methods. The genomic selection model was the best method compared to the single SNP, the single SNP with a random polygenic effect (K model), and the CAR score regression.

Recently, many instances of GWAS have been published with plant species, including barley, sorghum, wheat, rice, sugarcane, soybean and particularly maize (Ingvarsson and Street 2011). From the analysis of 271 inbreds genotyped for 28,626 single nucleotide polymorphisms (SNPs), Bernardo and Thompson (2016) calculated chromosomal effects from the effects of SNP alleles carried on the chromosomes. Many chromosome-inbred combinations showed large chromosome x inbred effects. Pace *et al.* (2015) carried out a GWAS with 384 maize inbred lines evaluated for 22 seedling root architecture traits and genotyped with 681,257 SNPs. They identified 268 marker-trait associations. Some of these SNPs were located within or near (less than one kilo base pairs) to candidate genes involved in root development at the seedling stage. Thirunavukkarasu *et al.* (2014) evaluated 240 elite inbred lines of subtropical maize under water stress and used a set of 29,619 high-quality SNPs. The GWAS identified 50 SNPs consistently associated with agronomic traits related to functional traits that could lead to drought tolerance. Thirty-one of the significant SNPs were situated near drought-tolerance genes. Schaefer and Bernardo (2013) used GWAS on a collection of 284 historical maize inbred lines and 39,166 SNPs and identified 19 QTLs for flowering time, 13 for kernel composition, and 22 for disease resistance. However, only two candidate genes were suggested: one regulating days to anthesis and one regulating oil concentration.

Genome-wide association studies with plant species have employed inbred lines panels. Thus, to our knowledge, no information is available on efficiency of GWAS in open-pollinated populations. Because association mapping has been primarily developed for mapping human genes, the available quantitative genetics theory, as that presented by Weir (2008), based on LD and analysis of case-control, is adequate for non-inbred populations. However, there is a lack of quantitative genetics theory for GWAS in inbred populations and inbred lines panels. In this paper we also evidenced the importance of fitting the additive-dominance model for traits showing unidirectional or bidirectional dominance and the additive model when there is no dominance, to achieve high power of QTL detection. We also highlighted the power of QTL detection, the false discovery rate (FDR), and the precision of GWAS. In summary, our objectives were to present additional quantitative genetics theory for GWAS, evaluate the relative efficiency of GWAS in non-inbred and inbred populations and in an inbred lines panel, and assess factors affecting GWAS, such as sample size and QTL heritability, effect of substitution, and dominance deviation. Importantly, the results for open-pollinated populations are directly applied to human populations.

## MATERIALS AND METHODS

The following theory aims to prove that a significant association between a SNP and a quantitative trait in an open-pollinated population, in a sample of recombinant inbred lines (RILs), and in an inbred lines panel, after correcting for population structure, depends on the LD between the SNP and at least one of the QTLs that affect the trait. Further, we present the parametric values of the average effect of a SNP substitution and LD measure in these association mapping populations, increasing the quantitative genetics knowledge on GWAS previously provided by human geneticists in the context of random population samples, case-control, and family-based studies.

### Quantitative genetics theory for GWAS in random cross populations

Consider a biallelic QTL (alleles **B**/**b**) and a SNP (alleles **C**/**c**) located in the same chromosome, and a population (generation 0) of a random cross species. Assuming LD, the joint gamete and joint genotype probabilities in the population are presented by Weir (2008). The QTL genotypic values are G_**BB**_ = m_b_ + a_b_, G_**Bb**_ = m_b_ + d_b_, and G_**bb**_ = m_b_ − a_b_, where m_b_ is the mean of the genotypic values of the homozygotes, a_b_ is the deviation between the genotypic value of the homozygote of higher expression and m_b_, and d_b_ is the dominance deviation (the deviation between the genotypic value of the heterozygote and m_b_). The average genotypic values of individuals with the SNP genotypes **CC**, **Cc**, and **cc** are

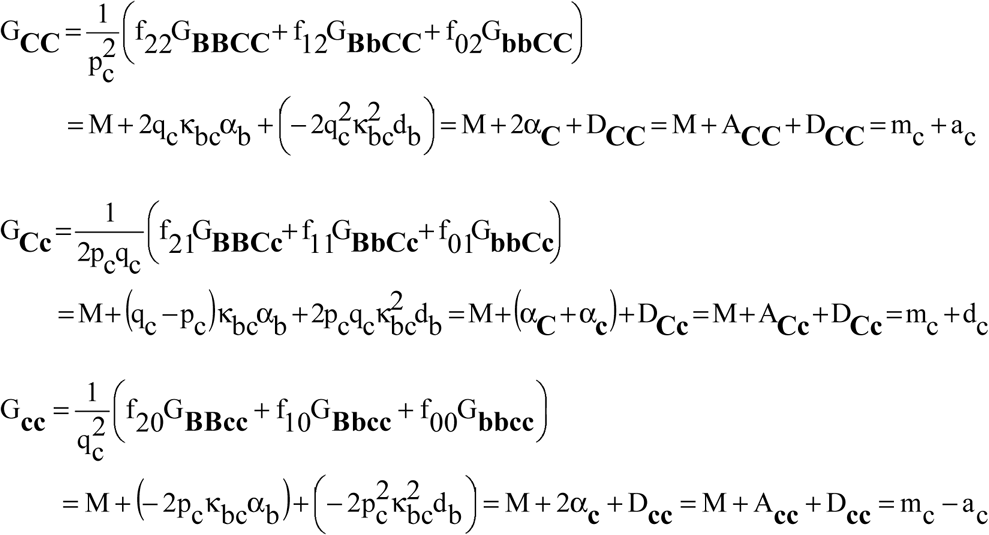

where p is the frequency of the major allele (**B** or **C**), q = 1 − p is the frequency of the minor allele (**b** or **c**), f_ij_ is the probability of the individual with i and j copies of the allele **B** of the QTL and the allele **C** of the SNP (i, j = 2, 1, or 0) (for simplicity, we omitted the superscript (0) - for generation 0 - in all parameters that depend on the LD measure of generation −1), 
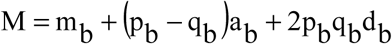
 is the population mean, 
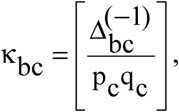

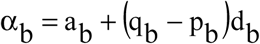
 is the average effect of a gene substitution, 
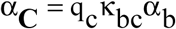
 and 
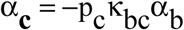
 are the average effects of the SNP alleles, and A and D are the SNP additive and dominance values. 
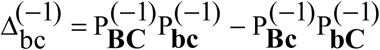
 is the measure of LD in the gametic pool of generation −1 (Kempthorne 1957), where P^(−1)^ indicates a joint gamete probability. Another common measure of LD is the square of the correlation between the values of the alleles at the two loci 
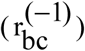
 in the gametic pool of generation −1 (Hill and Robertson 1968). Note that 
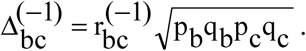
 The average effect of substituting the allele **C** for **c** is α_SNP_ = α_**C**_ − α_**c**_ = −κ_bc_α_b_. The dominance deviation for the SNP is 
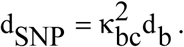
 The other SNP parameters are 
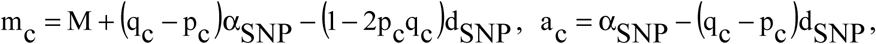
 and d_c_ = d_SNP_. The GWAS and genomic selection models commonly fit the SNP parameters a (values 1, 0, and −1 for SNP genotypes **CC**, **Cc**, and **cc**, respectively) and d (values 0, 1, and 0 for SNP genotypes **CC**, **Cc**, and **cc**, respectively).

Assuming no QTL in LD with the SNP 
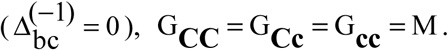
Thus, the identification of the QTL can be based on testing the hypothesis that there is no difference between these genotypic means. Assuming thousands of SNPs, it is necessary to employ a Bonferroni-typeprocedure to control the type I error when there are multiple-comparisons, as that proposed by Benjamini and Hochberg (1995). Note that 
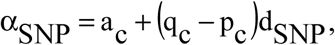
 where 
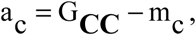

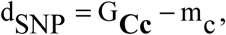
 and 
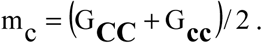

### Quantitative genetics theory for GWAS with inbred lines panel

In general, the inbred lines in a panel represent the genetic variability for the traits being assessed. Therefore, an inbred lines panel includes inbreds from distinct populations or heterotic groups. Consider again a QTL (alleles **B**/**b**) and a SNP (alleles **C**/**c**) located in the same chromosome, and that they are in LD in a population (generation 0). Assuming n (n → ∞) generations of selfing, the (limits of the) probabilities of the inbreds are (for simplicity, we omitted again the superscript (0) - for generation 0 - in all parameters that depend on the LD measure of generation −1)

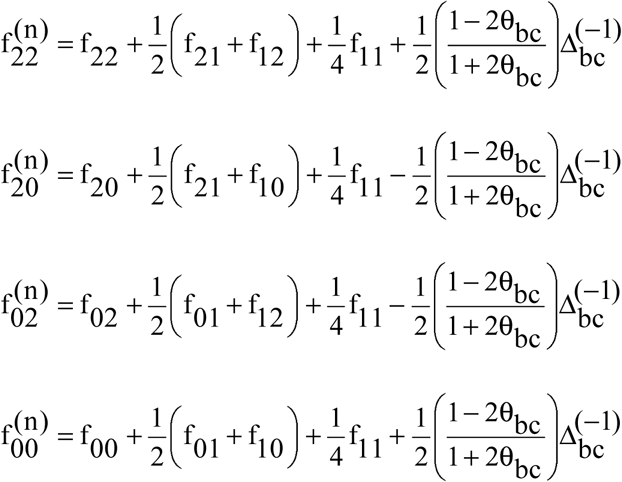

where θ_bc_ is the frequency of recombinant gametes. The haplotypes are 
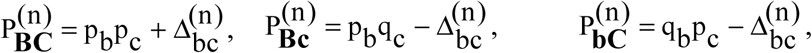
 and 
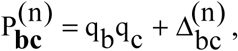
where 
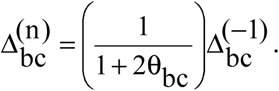
Thus, if there is crossing over (0 <θbc≤ 0.5), the LD in this inbredpopulation is lower than the LD in generation −1. If the SNP and QTL are completely linked (θ_bc_=0), the LD in the inbred population is the same LD in generation −1. The maximum decrease is 50%, achieved with θ_bc_ = 0.5. Compared with the LD in generation 0, the LD in generation n is 
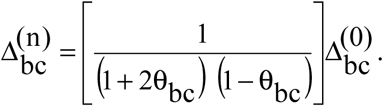
 Thus, the maximum decrease is approximately 10%, achieved with θ_bc_ = 0.25. In contrast, after n generations of random crosses 
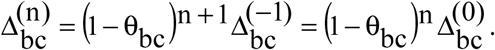
 Thus, if 0 < θ_bc_ ≤ 0.5, the maximum decrease is 100% ^since^

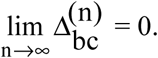

If the panel includes double haploid (DH) lines, the LD for the DH lines sampled from a population is 
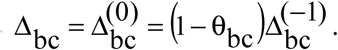
Thus, the LD in a sample of DH lines is greater than the LD in a sample of inbred lines (up to 12.5% greater when θ_bc_ = 0.25).

For the inbreds sampled from a population, we have

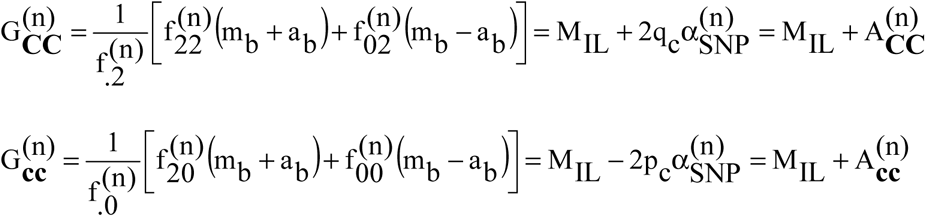
 where 
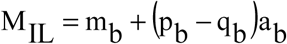
is the inbred population mean, 
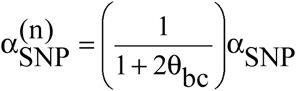
is the SNP average effect of allele substitution in the inbred population, and A is the SNP additive value for an inbred line. Assuming no QTL in LD with the SNP, 
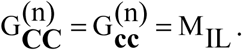
 Notice that
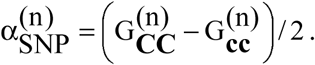
 In the case of DH lines, 
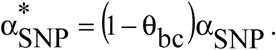
 Thus, assuming θ_bc_=0, the SNP average effect of substitution is approximately the same in a non-inbred and in an inbred population (RILs or DH lines).

The haplotypes of an inbred lines panel including inbreds from N populations are 
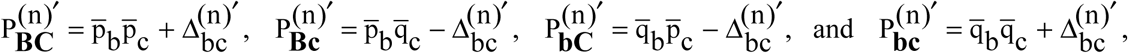
 where 
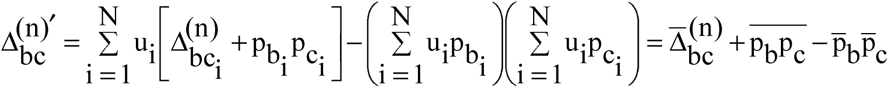
and u_i_ is the probability of an inbred line belonging to population i. Our simulated data showed that the LD value in an inbred lines panel tends to be lower than the LD in each group of inbreds (that are lower than the LD in the base populations) because it is an admixture of positive and negative LD values.

### Simulation

We simulated 50 samples of populations with LD using the software *REALbreeding* (Viana *et al.* 2017, 2016, 2013; Azevedo *et al.* 2015). This software has been developed by the first author using the program *REALbasic 2009*. Population 1, generation 0, is a composite of two populations in linkage equilibrium. Population 1, generations 10s and 10r10s, were obtained from Population 1, generation 0, assuming 10 generations of selfing and 10 generations of random crosses followed by 10 generations of selfing, respectively, assuming sample sizes of 100 and 400, respectively. Populations 2, 3, and 4, generation 10s, are also inbred populations (10 generations of selfing) derived from composites of two populations, also assuming a sample size of 100. The parents of populations 2 and 3 were assumed to be non-improved and improved populations, respectively. An improved population was defined as having frequencies of favorable genes greater than 0.5, while a non-improved population was defined as having frequencies less than 0.5. A composite is a Hardy-Weinberg equilibrium population with LD for only linked markers and genes. In the case of a composite of two populations in linkage equilibrium, 
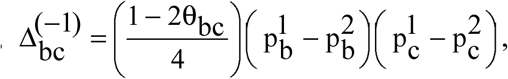
 where the indices 1 and 2 refer to the parental populations. Under random crosses, we instructed *REALbreeding* to generate two descendents by plant (one as male and one as female) and to allow selfing. Under selfing, *REALbreeding* used the single seed descent process. Thus, the individuals in generations 0 and the derived inbred lines are non-related and the individuals in generation 10r10s can be related.

Based on our input, *REALbreeding* randomly distributed 10,000 SNPs, 10 QTLs and 90 minor genes (QTLs of lower effect) in 10 chromosomes (1,000 SNPs and 10 genes by chromosome). The average SNP density was 0.1 cM. The genes were distributed in the regions covered by the SNPs. Four, three, two, and one QTLs were inserted in chromosomes 1, 5, 9, and 10, respectively. We also specified one SNP within each QTL (with same frequency) and a minimum distance between linked QTLs of 10 cM. To allow *REALbreeding* to compute the phenotypic value for each genotyped individual, we informed the minimum and maximum genotypic values for homozygotes, proportion between the parameter *a* for a QTL and the parameter *a* for a minor gene (a_QTL_/a_mg_), degree of dominance ((d/a)_i_, i = 1,…, 100), direction of dominance, and broad sense heritability. *REALbreeding* saves two main files, one with the marker genotypes and another with the additive, dominance, and phenotypic values (non-inbred populations) or the genotypic and phenotypic values (inbred populations). The true additive and dominance genetic values or genotypic values are computed from the population gene frequencies (random values), LD values, average effects of gene substitution or *a* deviations, and dominance deviations. The phenotypic values are computed from the true population mean, additive and dominance values or genotypic values, and from error effects sampled from a normal distribution. The error variance is computed from the broad sense heritability.

We simulated three popcorn traits. The minimum and maximum genotypic values of homozygotes for grain yield, expansion volume, and days to maturity were 30 and 180 g per plant, 15 and 65 mL/g, and 100 and 170 days, respectively. We defined positive dominance for grain yield (0 < (d/a)_i_ ≤ 1.2), bidirectional dominance for expansion volume (−1.2 ≤ (d/a)_i_ ≤ 1.2), and no dominance for days to maturity ((d/a)_i_ = 0). The broad sense heritabilities were 40 and 80%. These values can be associated with individual and progeny assessment, respectively. Assuming a_QTL_/a_mg_ = 10, the QTL heritabilities ranged from 3.2 to 11.8%. The GWAS was performed in population 1, generations 0 and 10r10s, and in the inbred lines panel obtained from inbreds of the populations 1 through 4, generation 0 (generations 10s). To assess the influence of the sample size on the GWAS efficiency, we considered sample sizes of 400 and 200. Thus, we used 100 or 50 inbreds from populations 1 through 4 to generate the inbred lines panel.

#### Statistical analyses

The analysis of the Q + K linear mixed model was performed with the software *GWASpoly* (Rosyara *et al.* 2016) fitting the additive and additive-dominance models for the open-pollinated population and the additive model for the RILs and inbred lines panel. For the population structure analysis, we used *Structure* software (Falush *et al.* 2003) and fitted the admixture model with correlated allelic frequencies and the no admixture model with independent allelic frequencies. The number of SNPs, sample size, burn-in period, and number of MCMC (Markov chain Monte Carlo) replications were 100 (10 random SNPs by chromosome), 400 (simulation 1), 10,000, and 40,000, respectively. The number of populations assumed (K) ranged from 1 to 7, and the most probable K value was determined based on the inferred plateau method (Viana *et al.* 2013). The population structure analysis evidenced four subpopulations (data not shown).

To classify each significant association as true or false, we used a program developed in *REALbasic 2009* by the first author. The classification criterion was based on the difference between the position of the SNP and the position of a true QTL (candidate gene). If the difference was less than or equal to 2.5 cM (Yu *et al.* 2008), the association was classified as true. The GWAS efficiency was assessed based on the power of QTL detection (probability of rejecting H_0_ when H_0_is false; control of the type II error), FDR (control of the type I error), and bias in the estimated QTL position (precision of mapping) (Li *et al.* 2010). We used Benjamini-Hochberg FDR of 5 and 1% to control the type I error (Benjamini and Hochberg 1995).

### Data availability

*REALbreeding* is available upon request. The data set is available at https://dx.doi.org/10.6084/m9.figshare.3201838.version3 Supplemental file S1 contains detailed description of all data files (SNP and QTL positions, SNP genotypes, and phenotypic values). Data citation:

Viana, José Marcelo Soriano; Mundim, Gabriel Borges; Pereira, Hélcio Duarte; Andrade, Andréa Carla Bastos; Fonseca e Silva, Fabyano (2017): Efficiency of genome-wide association study in random cross populations. figshare. https://dx.doi.org/10.6084/m9.figshare.3201838.version3

## RESULTS

Our first result from the open-pollinated population, assuming sample size of 400 and FDR of 1%, was disappointing since most grain yield QTLs of high heritability showed low power of detection. For example, the power of detection for the QTLs with heritabilities of 8.4, 9.4, and 11.6% were 4.2, 8.3, and 10.4%, respectively (Figure 1a). We realized that the problem was the high dominance deviation for these QTLs (the greatest values among the 10 QTLs). This explained the relatively low coefficients of determination for the linear regression models relating QTL detection power and heritability, especially with sample size of 200 and FDR of 5% (45, 55, and 19%) (Figure1a, c, and Figure 2a). The solution to this problem was to fit the additive-dominance model for grain yield and expansion volume. Regardless of the sample size and FDR, for 76% of the grain yield and expansion volume QTLs the detection power was increased (Figures 1b, d, and Figure 2b). Previously undetected QTLs showed detection power ranging from 2.3 to 52.3%. The increase in the detection power for previously detected QTLs ranged from 0.8 to 2,300% (244.6% on average), mainly with sample size of 400 and FDR of 1%. A consequence of fitting the additive-dominance model for grain yield and expansion volume was an increase in the coefficients of determination for the linear regression models relating QTL detection power and heritability, especially with lower sample size (R^2^ of 79, 81, and 46%) (Figure1b, d, and Figure 2b). Unlike, the additive-dominance model should not be fitted for traits showing no dominance. Fitting the additive-dominance model for days to maturity, for 82% of the QTLs the detection power decreased from 6.6 to 100.0% (48.4% on average), regardless of sample size and FDR.

**Figure 1.**
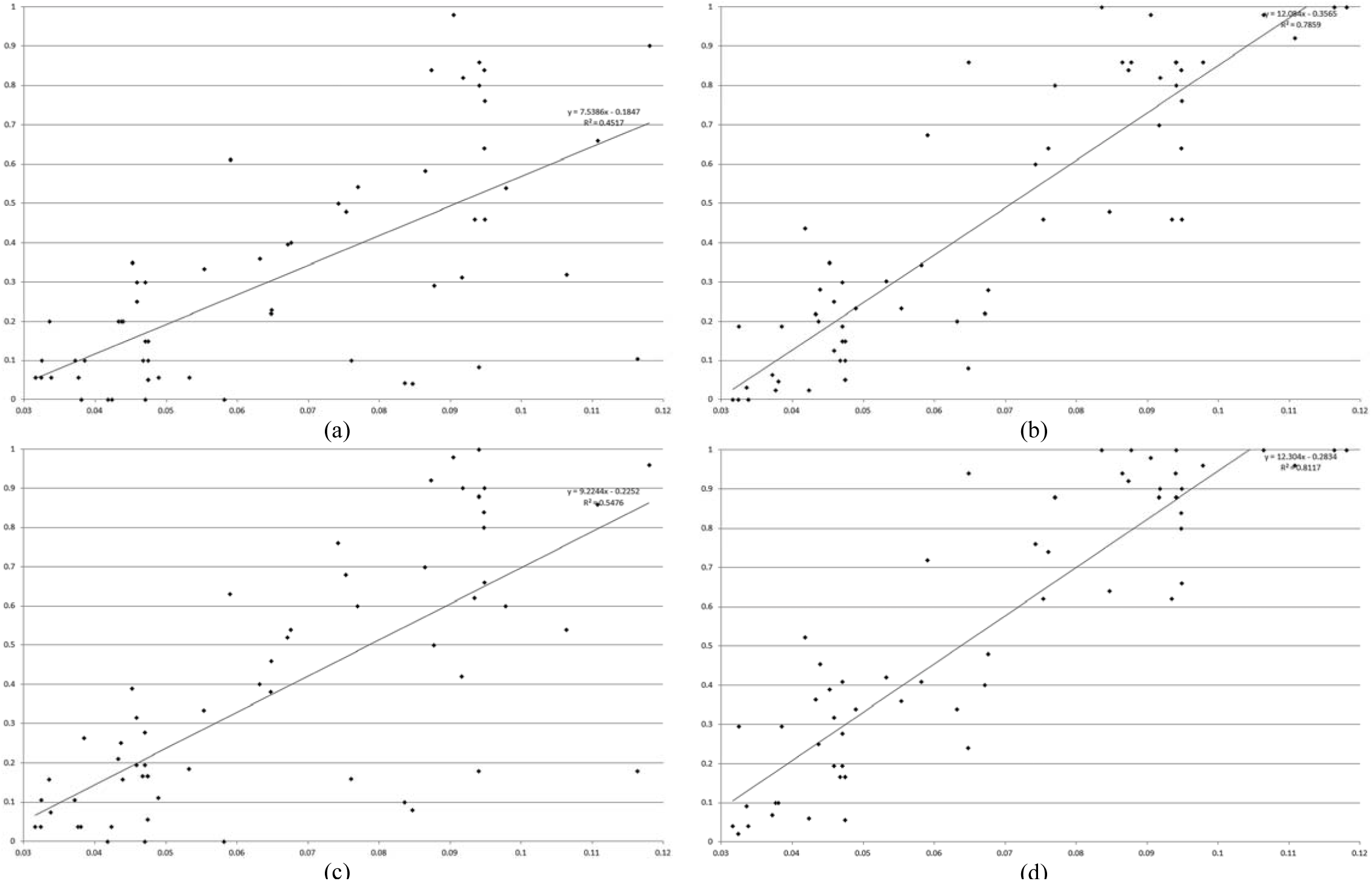
Relationship between QTL heritability (X axe) and power of detection (Y axe) concerning population 1, generation 0, and QTLs determining grain yield, expansion volume, and days to maturity, assuming additive (a, c) and additive-dominance models (b, d), sample size of 400, and levels of significance of 1 (a, b) and 5% (c, d).

Defining low, intermediate, and high heritability QTLs as those with heritability values less than or equal to 3.5%, between 3.6 and 7.5%, and greater than 7.5%, respectively, the power of detection was maximized increasing the sample size from 200 to 400 and the FDR from 1% to 5% (Table 1). It is important to highlight that under sample size of 400 the observed FDR was equal to or lower than the specified level of significance. Assuming sample size of 400 and FDR of 5%, the average power of detection for the low, intermediate, and high heritability QTLs were 9.7, 32.7, and 87.7%, respectively. The minimum and maximum values were 2 and 29.5%, 5.5 and 94%, and 62.0 and 100.0%. The observed FDR was 3.8%. Decreasing the sample size to 200 decreased the QTL detection power (28, 67 and 62% for the low, intermediate and high heritability QTLs, respectively) and increased the observed FDR to 9.4%. Concerning the bias in the QTL (candidate gene) position, it should be also highlighted that at least 97% (assuming sample size of 200 and FDR of 5%) of the QTLs were declared by the SNP within it, and that the number of significant SNPs within the range of 2.5 cM was very low, regardless of sample size and FDR. The average number of significant SNPs within the range of 2.5 cM varied from 0.1 to 1.0. The average bias in the QTL position from a significant SNP within the range of 2.5 cM varied from approximately 0.2 to 0.4 cM. Furthermore, we also observed that the QTL detection power has low correlation with the QTL average effect of substitution (0.1) and QTL dominance deviation (0.2). The correlation between QTL detection power and QTL heritability ranged from 0.68 to 0.90, proportional to the sample size.

**Table I.**
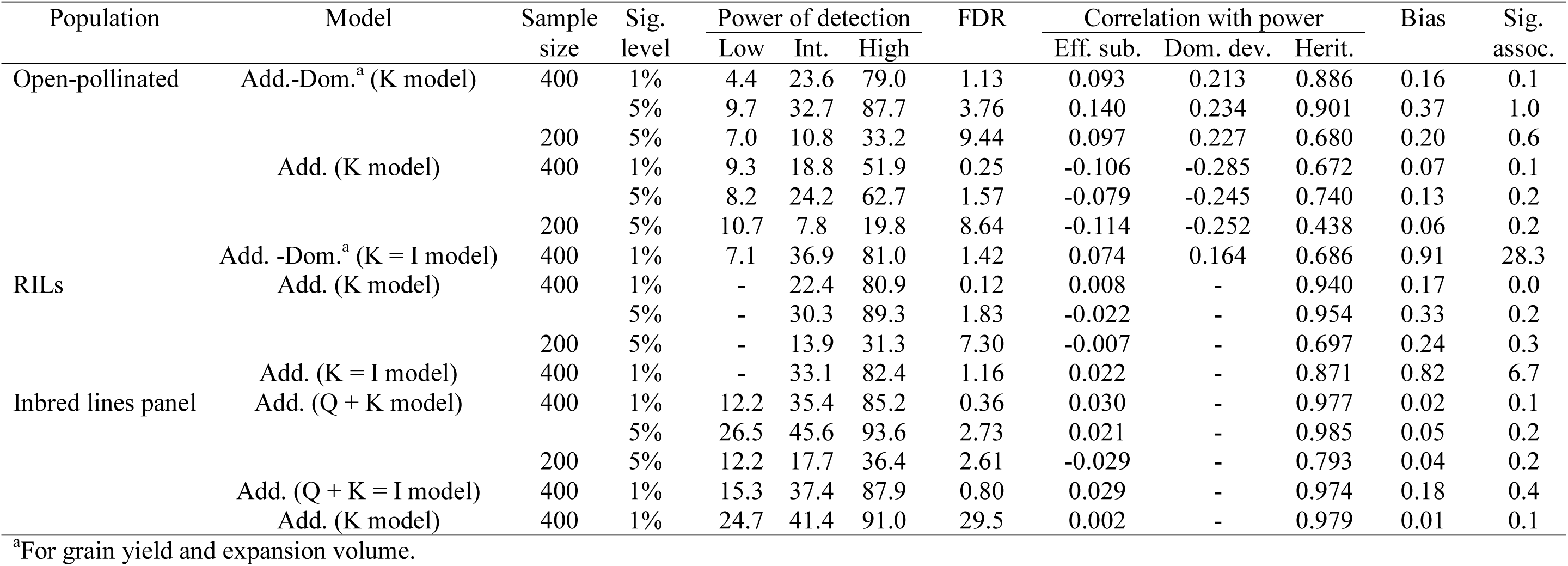
Average power of detection (%) for the low (≤ 3.5%), intermediate (3.6-7.5%), and high (≥ 7.6%) heritability QTLs, false discovery rate (%), correlations between power and QTL effect of substitution, dominance deviation, and heritability, bias in the QTL position for significant associations outside the QTL (cM), and number of significant SNPs outside the 2.5 cM interval, regarding population 1, generations 0 (open-pollinated) and 10r10s (RILs), and an inbred lines panel, five models, two sample sizes, and two levels of significance.

The analysis of the RILs from the open-pollinated population provided essentially the same results (similar magnitude of the statistics) concerning QTL detection power, control of the type I error, and mapping precision (Figure 2c, d, and Table 1). The only significant difference was the absence of low heritability QTLs. We can highlight a slightly better control of the type I error also.

**Figure 2.**
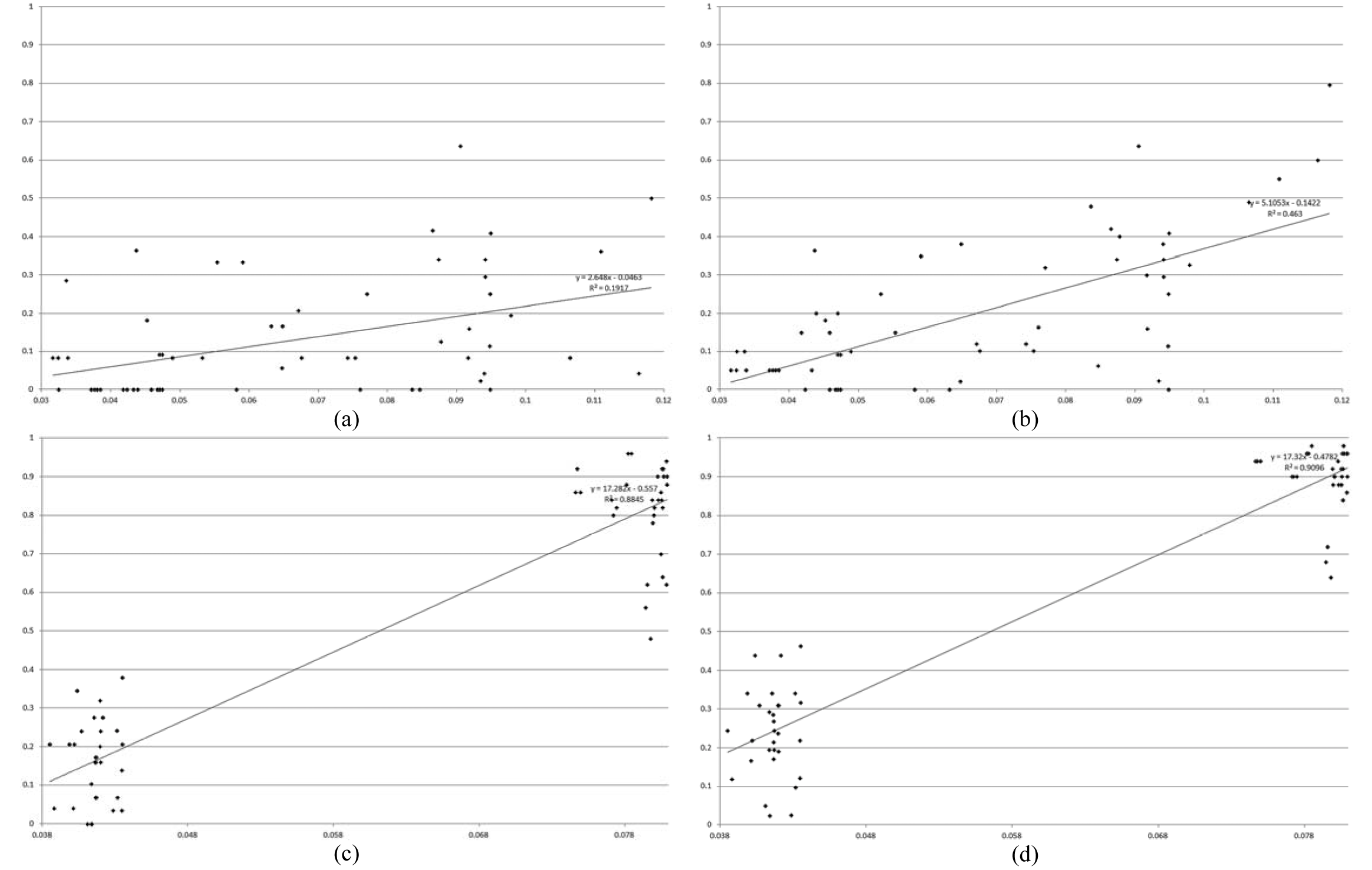
Relationship between QTL heritability (X axe) and power of detection (Y axe) concerning population 1, generations 0 (a, b) and 10r10s (c, d), and QTLs determining grain yield, expansion volume, and days to maturity, assuming additive (a, c, d) and additive-dominance models (b), sample sizes of 400 (c, d) and 200 (a, b), and levels of significance of 1 (c) and 5% (a, b, d).

Thus, the QTL detection power was also maximized assuming sample size of 400 and FDR of 5% and the observed FDR was 1.8% in this scenario. The decrease in the sample size significantly decreased the power of QTL detection (54 and 65% for the intermediate and high heritability QTLs, respectively) and increased the FDR (to 7.3%) too. Regarding of the inbred lines panel, it is impressive to realize that this population provided an improvement in the outstanding results offered by the non-inbred and inbred open-pollinated populations, concerning low and intermediate heritability QTL detection power, control of the type I error, and mapping precision (Figure 3a, b, and Table 1). The increase in the low heritability QTL detection power ranged from 74 to 177%. For the intermediate heritability QTLs the increase ranged from 39 to 64%. The observed FDR was reduced in up to 72% and the bias in the QTL position decreased between 80 and 87%. The QTL detection power was also maximized assuming sample size of 400 and FDR of 5%, combined with an observed FDR of 2.7%. Similarly to non-inbred and inbred open-pollinated populations, the decrease in the sample size significantly decreased the power of QTL detection (54, 61 and 61% for the low, intermediate, and high heritability QTLs, respectively) but the FDR was unaffected.

**Figure 3.**
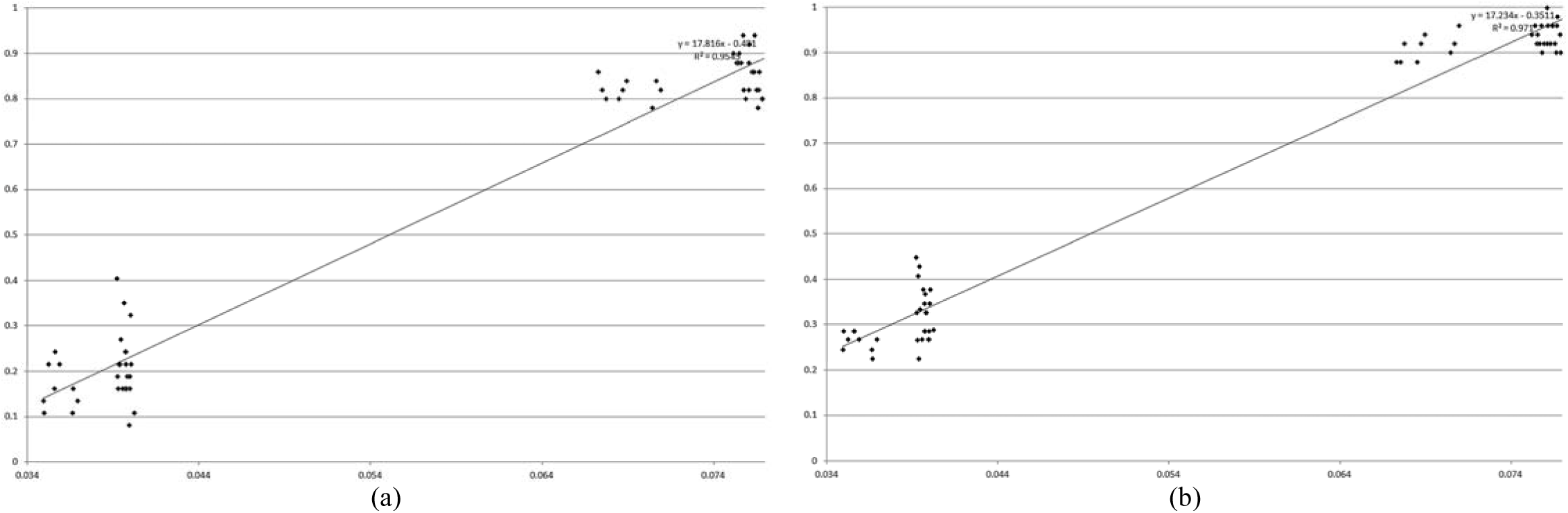
Relationship between QTL heritability (X axe) and power of detection (Y axe) concerning the inbred lines panel and QTLs determining grain yield, expansion volume, and days to maturity, assuming sample size of 400 and levels of significance of 1 (a) and 5% (b).

Finally, it is important to highlight that the FDR is mainly affected by population structure, compared to relationship information. Assuming non-inbred population, sample size of 400 and FDR of 1%, ignoring the relationship information (by ignoring the polygenic effect), the observed FDR was unaffected (1.1 vs. 1.4%) but the number of significant associations outside the 2.5 cM interval was drastically increased (from practically zero to 28; Table 1). As will be discussed further, these are not all false-positive associations but due to LD between the SNPs and one or more QTLs in the chromosome. With RILs, because the level of LD is lower than in the non-inbred population, the observed FDR was 1.2% but the number of significant associations outside the 2.5 cM interval achieved approximately 7. For inbred lines panel, ignoring only the relationship information the FDR was 0.8% but ignoring only population structure increased the FDR to 29.5% Ignoring the relationship information and population structure in the analysis of the inbred lines panel determined thousands of significant associations in all chromosomes, increasing drastically the FDR (to approximately 60%; data not shown). In these scenarios it was not possible to detect QTLs because many significant associations were observed along the length of a chromosome or in one or more large chromosome regions, especially for chromosomes 1 (four QTLs) and 5 (three QTLs) (data not shown).

## DISCUSSION

The presented theory proves that a significant association from a GWAS in a non-inbred or inbred random cross population and in an inbred lines panel, correcting for population structure and relatedness, is due to LD between the SNP and one or more linked QTLs. The theory also shows that GWAS provides estimation of the average effect of a SNP substitution (and consequently the estimation of SNP effects). Using SNP effects to measure chromosome and chromosome x inbred effects, Bernardo and Thompson (2016) showed that GWAS also provide dissection of the germplasm architecture for quantitative traits. Schaefer and Bernardo (2013) estimated SNP effects and identified candidate genes and QTL hot spots (chromosome regions with previously mapped QTLs) for days to flowering, kernel composition, and disease resistance. Based on the theory presented, only if there is a single QTL in LD with a significant SNP, if the SNP is within the QTL, and if QTL and SNP alleles have the same frequency it is adequate to consider the SNP average effect of substitution as the QTL average effect of substitution. We additionally provided the parametric values of SNP effects commonly fitted in the GWAS and genomic selection models, and the genotype and gametic probabilities and the parametric LD values in a completely inbred population and in an inbred lines panel. The LD in a group of RILs are lower than the LD in the non-inbred population and the LD value in an inbred lines panel tends to be lower than the LD in each group of inbreds (that are lower than the LD in the base populations) because it is an admixture of positive and negative LD values.

To our knowledge, this is the first study on GWAS efficiency in open-pollinated population. The results are impressive and show that the identification of candidate genes can be highly efficient, depending on sample size and QTL heritability. LD is for sure another important factor affecting GWAS (Weir 2008). Thus, based on a sample of 400 individuals and defining a level of significance of 5%, the power of detection of low (≤ 3.5%), intermediate (3.6−7.5%), and high (≥ 7.6) heritability QTLs can achieve approximately 30, 90, and 100%, respectively (10, 30, and 90% on average). This result is achieved keeping the FDR bellow 5% and is associated with a very low number of significant associations close to the QTL (highly precise mapping), besides the significant SNP within the QTL. This means that GWAS efficiency is maximized when there is at least one SNP within each QTL, with the same allelic frequency. This seems very restrictive and, unfortunately, is. To achieve high efficiency when there is not a SNP within each QTL, high LD between a SNP close to the QTL and greater sample size are required, especially for low heritability QTLs. In a random cross population the LD measure depends also on the SNP and QTL allele frequencies. Thus, significant associations involving few SNPs with the same QTL can be observed, including SNPs that are tens of mega base pairs (or centiMorgans) from the QTL. In reality, a closely linked QTL and SNP can have a lower LD value compared to a more distant QTL and SNP pair. In populations with low level of LD, significant associations are expected to occur for only SNPs within the QTL or located very close to the QTL (within a few hundred base pairs), which favors the identification of a candidate gene for the QTL. In this scenario, a QTL would be declared based on one to a small number of significant associations spanning a chromosome region of a few kilo base pairs (not mega base pairs or centiMorgans).

Field results have demonstrated that GWAS are best carried out with a large sample size (Yu and Buckler 2006). According to Flint-Garcia *et al.* (2005), increasing the population size increases the number of individuals with rare alleles, thus improving the power to test the association between these rare alleles and the trait of interest. Yu *et al.* (2008) showed that the gain in the GWAS efficiency by increasing sample size was evidenced by increased power of QTL detection and smaller FDR, mainly with heritability of 0.7 in comparison with a heritability of 0.4. Based on a simulation study, Long and Langley (1999) demonstrated that approximately 500 individuals should be genotyped for 20 SNP loci within the candidate gene region to detect marker-trait associations for QTLs that account for as little as 5% of the phenotypic variation. They observed that more power was achieved by increasing the population size than by increasing the SNP density within the candidate gene region.

Our most significant contribution to the knowledge on GWAS is the empirical proof that the additive-dominance model must be fitted for traits controlled by dominance (uni- or bidirectional), to increase the QTL power of detection. We additionally evidenced that the additive-dominance model must not be fitted for traits determined only by additive gene effects, to avoid a decrease in the QTL detection power. This is probably due to over fitting. Further, we provided results for comparing GWAS in non-inbred and inbred random cross populations, and in an inbred lines panel. The inbreeding did not affect the GWAS efficiency, but RILs, if available, can be interesting to maximize the QTL heritabilities, since they allow standard experimental procedures (local control and replication) and the assessment of SNP x environment interaction. Compared to GWAS in an inbred lines panel, GWAS in random cross population was less efficient, i.e., showed lower power of QTL detection for the low and intermediate heritability QTLs, slightly inferior control of false-positive associations, and higher bias in QTL position. This is due to higher genetic variability in the inbred lines panel since the average LD in the population 1, generation 0, is higher than the average LD in the inbred lines panel (average absolute Δ equal to 0.0403 and 0.0249, respectively). The genetic variability in the inbred lines panel is 9 to 13 times greater, depending on the trait (data not shown).

According to Flint-Garcia *et al.* (2005), the inbred lines panel exploits the rapid breakdown of LD in diverse maize lines, enabling very high resolution for QTL mapping. Population structure results from constructing a panel with inbreds from various breeding programs and distinct heterotic groups, which can cause false-positive marker-trait associations if the data is not corrected (Yan *et al.* 2009). The lowest parametric LD values for the inbred lines panel occurred in published studies (Yan *et al.* 2009, Remington *et al.* 2001). Moreover, with the inbred lines panel, generally, only SNP loci within the QTL showed significant association, which is a highlighted result from GWAS that can serve as a basis for a fine mapping strategy for marker-assisted selection and map-based cloning genes (Gupta *et al.* 2005).

Our results are comparable to previous GWAS with field and simulated data. Concerning the QK model, Bernardo (2013) observed that the power of QTL detection and number of false-positive associations were proportional to the sample size. Assuming FDR of approximately 1% and an average QTL heritability of approximately 5%, the power of detection increased from 13 to 45% when the sample size increased from 384 to 1,536. In the study of Yang *et al.* (2010) the QTL detection power was relatively low for QTLs with heritability lower than 10% but increased significantly with the increase in the population size. Assuming sample size of 155, the power of detection was 16.5, 59.2, and 87.6% for the low (1%), intermediate (5%) and high (≥ 10%) heritability QTLs, respectively. Yu *et al.* (2008) investigated the genetic and statistical properties (power of QTL detection and FDR) of the nested association mapping (NAM) design. With 5,000 genotypes, they achieved an average power of QTL detection of 57% (with a range of 30 to 85%) when considering two trait heritabilities (0.4 and 0.7) and two different numbers of QTL controlling the trait (20 and 50). They also observed that a higher heritability always gave higher QTL detection power, particularly for QTL with moderate to small effect. However, the FDR values were high, ranging from 9 to 23%.

Concerning the relevance of relatedness, even if due to identity by state, and population structure correction, our findings agreed with previous knowledge that the best GWAS model must include a polygenic effect - to eliminate significant associations outside of the QTL interval (including false-positive associations) - and a population structure effect - to control the type I error, as highlighted by Yu et al (2006) and Bernardo (2013), among others. Stich and Melchinger (2009) and Yang *et al.* (2010) observed best control of spurious associations by the K model. In the study of Flint-Garcia (2005) the population structure effect was significant, explaining 9.3% of the phenotypic variation, on average.

The GWAS in plant breeding has been effective for identifying candidate genes for quantitative traits such as plant architecture, kernel composition, root development, flowering time, drought tolerance, pathogen resistance, and metabolic processes (Zhu *et al.* 2008). Our study provided the following additional knowledge: 1) the additive-dominance model must be fitted for traits controlled by dominance effects but must not be fitted for traits controlled only by additive effects, to achieve high power of QTL detection; 2) with sample size of 400 and level of significance of 5%, the power of detection for the low, intermediate, and high heritability QTLs can achieve approximately 30, 90, and 100%, respectively; 3) under sample size of 400, the observed FDR was equal to or lower than the specified level of significance; 4) GWAS in random cross populations is highly precise, since at least 97% of the QTLs were detected by the SNP inside it and the number of significant associations outside of the QTL interval (2.5 cM) is very low; 5) inbreeding does not affect the GWAS efficiency; 6) identity by state is important to control significant associations outside of the QTL interval; and 7) in random cross populations, FDR is mainly affected by population structure, compared to relationship information. Based on our evidence, breeders can employ non-inbred and inbred populations for GWAS while taking into account that the level of LD should be high, the sample size should be higher than that necessary for QTL mapping, and the QTL heritability should be intermediate to high to achieve greater power of QTL detection and precise mapping of candidate genes.

## ACKNOWLEDGMENTS

We thank the National Council for Scientific and Technological Development (CNPq), the Brazilian Federal Agency for Support and Evaluation of Graduate Education (Capes), and the Foundation for Research Support of Minas Gerais State (Fapemig) for financial support.

